# Creativity drives performer–listener emotional and physiological alignment in live music improvisation

**DOI:** 10.64898/2026.05.27.726846

**Authors:** Ioanna Zioga, Clément Guichet, Caroline Di Bernardi Luft, Charalabos Papageorgiou, Christina Anagnostopoulou

## Abstract

Music is a fundamental medium for human communication, yet it remains unclear whether it enables genuine alignment of internally felt emotions beyond mere emotion perception. Creative expression may be central to this process: when performers improvise with creative intent, they engage in self-expression that could draw listeners into closer emotional and physiological resonance. Here, dual-electrocardiography was recorded across 37 performer-listener dyads while performers generated live improvisations under conventional, unconventional, or creative instructions. We show that creative improvisation enhances dyadic emotional alignment for both valence and arousal, and increases cardiac synchronization. Performer-listener alignment was strongest when performers’ felt and expressed emotions were themselves aligned, and when listeners showed a gradual arousal buildup. Creativity also elicited higher sublimity, lower unease and vitality, and greater diversity of selected emotional labels. These findings establish creativity as a mechanism through which music achieves one of its most celebrated functions: the creation of shared human experience.

## 1. Introduction

Music is frequently described as a universal communication medium, capable of evoking shared emotional experiences even across disparate linguistic and cultural backgrounds^1^. Traditionally, music research has examined how listeners perceive emotions expressed by the music^2^. However, it remains unclear whether this transmission reflects a genuine coupling of the performer’s internal emotional experience with that of the listener. In this study, we investigate how creativity facilitates this emotional alignment between performer-listener dyads during live music improvisation.

A critical distinction exists between the emotions expressed by the music and the emotions actually felt by each individual^3,4^. For instance, a listener might experience joy while recognizing that a musical piece expresses melancholy. Meta-analytic evidence suggests that these expressed emotions are typically more intense than the resulting felt emotions^3,5^. While the majority of previous research has focused on either the structural features that signal expressed emotion or the subjective felt emotion of the listener in isolation, we shift the focus toward the relationship between the performer’s and the listener’s felt emotions.

Here, we define this relationship, dyadic emotional alignment, as the degree of convergence between performer-listener self-reported felt emotions during a shared musical experience. Importantly, this construct differs from emotion communication accuracy, which concerns whether listeners correctly infer the emotion expressed by the performer or the music^6^. Dyadic emotional alignment instead captures the extent to which both individuals subjectively experience similar affective states during the interaction.

Recent research has increasingly utilized live performance settings, both in the lab and in situ, to investigate the behavioural and neurophysiological foundations of musical experience^7–10^. The physical presence of a performer has been shown to heighten audience engagement and facilitate heart rate synchronization between audience members^8,10^, particularly when the music is familiar^7^. Furthermore, live performance enhances neural phase-locking to musical rhythm^11^, reinforcing the view that music is inherently social^12,13^. This social dimension is critical, as sharing musical experiences can amplify the resulting pleasure^14^. Importantly, the nature of the social bond plays a role in these dynamics; interpersonal familiarity is associated with distinct patterns of inter-brain synchronization, often resulting in higher synchrony among friends compared to strangers^15,16^.

Previous studies have also shown that the neurophysiological signals of audiences synchronize during listening^17–21^. Evidence that a more creative approach changes interpersonal coupling comes from Nozawa and colleagues^22^, who contrasted a “Let-go” (more improvisatory) performance mode with a “Strict” mode in a concert experiment. They found that the creative approach affects movement synchrony in the audience in opposite directions, depending on the timescale: Let-go performances reduced synchrony compared to Strict in shorter timescales, while they enhanced synchrony on longer timescales.

Furthermore, listeners’ arousal often unfolds as a trajectory over time and can show build-ups toward climaxes^5,23,24^. Indeed, sustained engagement with unfolding musical structure may contribute not only to subjective emotional experience, but also to interpersonal alignment between listeners and performers. Interestingly, specific modes of engagement, such as “giving oneself over to the music”, have been associated with lower physiological synchrony, likely reflecting an increase in inter-individual variability during deeply subjective experiences^10^. Crucially, no study to date has explored whether performer-listener synchronization is modulated by creative intent or whether dynamic changes in listener engagement contribute to performer-listener emotional alignment. The present study addresses this gap by examining how the instruction to be creative shapes emotional alignment and physiological coordination within performer-listener dyads.

Creativity is a multifaceted cognitive capacity, often conceptualized as the novel combination of existing elements. While definitions vary, a broad consensus characterizes creativity as the production of ideas that are both original and appropriate within a given context^25,26^. Despite a growing body of research into musical creativity, few studies have operationalized the construct as a multidimensional variable in controlled experimental settings. Here, we operationalize creativity through a tripartite framework, instructing performers to improvise under conventional, unconventional, or creative conditions. Research in the verbal domain has consistently shown that explicit instruction to “be creative” enhances the originality of outputs^27–31^. For instance, prompted creativity leads to the generation of words with greater semantic distance compared to prompts for unconventional (novel but inappropriate) or conventional (appropriate but not novel) responses^32^.

Creative musical experiences differ not only in the intensity of emotional alignment they produce, but also in the qualitative nature of the emotions evoked^33^. Music-induced emotions often extend beyond basic valence and arousal dimensions to include aesthetic states such as wonder, transcendence, peacefulness, and sublimity^33–35^. In parallel, neurocognitive research suggests that creative generation relies on broader semantic activation and access to more remote conceptual associations^36–39^. Consequently, creative improvisations may prompt more diverse emotional descriptions, manifested as greater lexical richness in participants’ emotional descriptions.

In the present study, we implemented a novel experimental paradigm pairing an expert improviser with a listener. The performer was tasked with generating original musical improvisations with one of three specific instructions: *conventional, unconventional, or creative*, while the cardiac activity of both individuals was recorded concurrently. Following each performance, both members of the dyad reported their felt emotions as well as the emotions they perceived the music to express. This paradigm was designed to address two primary objectives: first, disentangle felt versus expressed emotional alignment within the dyad; and second, isolate the specific role of creativity in emotion transmission by manipulating its distinct facets through explicit instruction. Musical improvisation provides an ideal model for studying real-time creativity and emotion. Each performance represents a unique creation that inherently eliminates familiarity biases, thus allowing for the experimental manipulation of creative intent.

We formulated two sets of hypotheses. *First*, we predicted that creative improvisations would enhance dyadic emotional alignment, and that this convergence would be reflected at the physiological level through increased cardiac synchrony. We further hypothesized that these effects would be moderated by interpersonal familiarity, and that dyadic alignment would be positively associated with listener engagement indexed by a gradual buildup of arousal over time. *Second*, we examined whether creativity alters the qualitative nature of emotional experience. Specifically, we tested whether creative improvisations elicit distinct aesthetic emotions and more diverse emotional descriptions, reflected in increased lexical richness and semantic variability across emotion categories.

## 2. Methods

### 2.1 Participants

Thirty-seven performer-listener dyads of healthy adults (*N* = 74) participated in the study. The final number of dyads involved in the present study exceeded the a priori sample size (*n* = 28) estimated by means of a statistical power analysis using G*Power^40^ (evaluated for 1-*β* = 0.80, *α* = 0.05, and effect size *f* = 0.25 for a repeated-measures within-factors ANOVA). This larger sample size was recruited to maintain robust statistical power against the low signal-to-noise ratio and risk of data loss typical of physiological recordings in live performance settings.

Of these, 11 dyads were friends (10 female, 12 male), while 26 were strangers (24 female, 28 male). Performers were required to have at least five years of musical training and be comfortable with improvisation (11 female; age range = 19-64 years, *M* ± *SD* = 39.78 ± 10.91). Independent *t*-tests confirmed significant differences between the two groups across expertise metrics. Specifically, performers demonstrated significantly higher musical training scores on the Gold-MSI Musical Training subscale^41^ compared to listeners (performers: *M* ± *SD* = 41.35 ± 4.28; listeners: *M* ± *SD* = 30.28 ± 12.95); *t*(44.3) = 4.88, *p* < 0.001, Cohen’s *d* = 1.15). Furthermore, a similar difference was observed in improvisation experience (performers: *M* ± *SD* = 9.15 ± 3.96 years; listeners: *M* ± *SD* = 2.79 ± 4.05 years; *t*(72) = 6.78, *p* < 0.001, Cohen’s *d* = 1.59). The most represented instrument was the piano (*N* = 10), followed by voice (*N* = 7) and electric guitar (*N* = 5). Additional instruments included saxophone (*N* = 4), classical guitar (*N* = 3), electric bass (*N* = 2), and clarinet (*N* = 2), with single participants playing violin, double bass, flute, and traditional instruments (lute, bouzouki, lyra). Listeners reported no hearing problems (*N* = 37; 22 female; age range = 18-60 years, *M* ± *SD* = 33.31 ± 12.51).

All participants provided written informed consent prior to participation. The study was approved by the Ethics and Deontology Research Board of the National and Kapodistrian University of Athens (approval no. 218/2025) and was conducted in accordance with the Declaration of Helsinki.

### 2.2 Experimental design & procedure

Participants arrived in pairs and sat facing each other at a distance of approximately four meters. They first completed the Goldsmiths Musical Sophistication Index (Gold-MSI)^41^ Musical Training scale to assess musical expertise and a question on years of improvisation experience. More details about the questionnaires can be found in Supplementary Note 1.

The experimental task consisted of a series of live musical improvisation trials, each followed by an emotional assessment (Figure 1). Before each trial, the performer received a prompt on a computer screen, unseen by the listener, instructing them to perform according to one of three styles:

**Figure 1.**
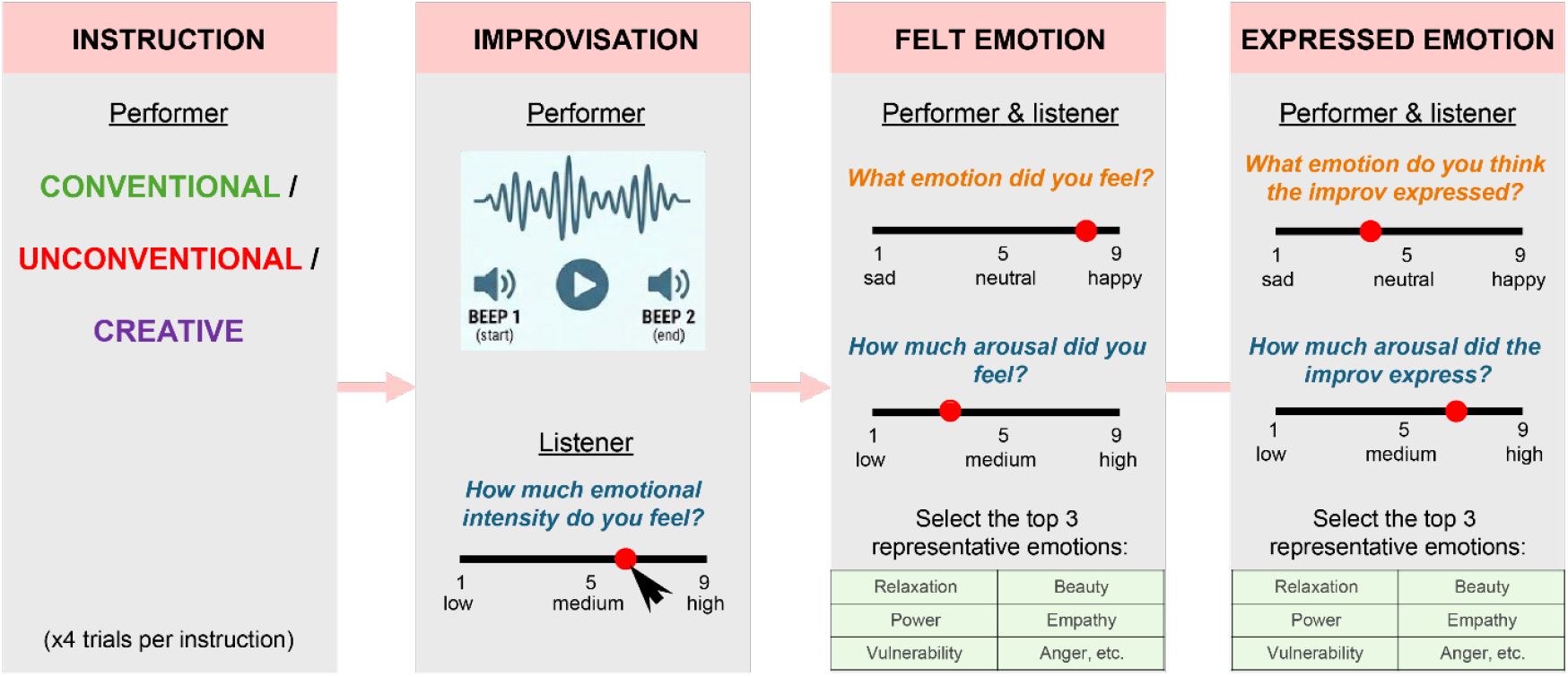
Illustration of the experimental design. Performers receive one of three randomized instruction prompts (conventional, creative, or unconventional) to guide a 90-s live improvisation. Listeners, blinded to the instruction, provide continuous behavioural ratings of their arousal using a digital slider interface. Following each improvisation, both members of the dyad complete an emotion assessment across two distinct axes: their internally felt emotions and their perceived emotional expression of the music. This assessment comprises Likert-scale ratings for valence and arousal, alongside the selection of specific aesthetic, music-related emotions from a predefined list.

1. Conventional: play something common, familiar, and predictable
2. Unconventional: play something unpredictable, unorthodox, experimental
3. Creative: play something imaginative, original, and fluent.

Each improvisation trial lasted 90 s, with the onset and offset marked by auditory beeps; participants were blinded to the trial duration. The experimental session consisted of 12 pseudorandomized trials (4 conventional, 4 unconventional, and 4 creative). Throughout each performance, the listener’s screen displayed the prompt: “How much emotional intensity do you feel?”, and listeners provided continuous, real-time behavioural ratings of their experienced arousal using a digital slider on a 9-point scale (1 = low, 5 = medium, 9 = high).

Following each trial, both the performer and the listener completed a comprehensive emotional assessment. This was divided into two distinct perspectives: internally felt emotions, and perceived emotional expression (the emotions they believed the music expressed). For both perspectives, participants provided responses on their respective computers across two distinct formats:

1. Likert ratings: Scoring of valence and arousal on a 9-point Likert scale, from 1 (very negative/low arousal) to 9 (very positive/high arousal),
2. List selection: Selection of the three most representative emotions from a predefined list of 31 music-related emotions. The list was compiled by adapting and extending the Geneva Emotional Music Scale^33^ (see Supplementary Note 2 for full details and label mapping).

#### Expert validation of instruction conditions

A validation was conducted to verify performer compliance with the instruction conditions. An independent pool of 7 expert musicians (7 males; age range = 19-45 years, *M* ± *SD* = 38.00 ± 9.09) evaluated the recorded improvisations. All experts possessed extensive musical training (Gold-MSI: *M* ± *SD* = 43.33 ± 2.80) and experience in musical improvisation (*M* ± *SD* = 9.00 ± 3.74 years). Experts were blinded to the experimental instructions under which the performances were generated. Stimuli were rated on conventionality, unconventionality, and creativity using 7-point Likert scales (1 = not at all, 7 = very much). Judges used an interactive audio player that allowed free navigation within each excerpt. The task was programmed using PsychoPy^42^. The study was deployed online and hosted on Pavlovia (https://pavlovia.org).

### 2.3 Signal data acquisition

#### Physiological recording

Physiological signals were recorded concurrently from the listener and the performer. Wireless Callibri sensors (Biotech, Russia; sampling rate: 1 kHz) were used to acquire electrocardiogram (ECG) data from both participants. A single-lead bipolar montage was implemented, with the sensor positioned vertically along the mid-sternum.

#### Setup and synchronization

The hyperscanning setup consisted of two laptops synchronized via a crossover Ethernet cable. All data streams were managed via the Lab Streaming Layer (LSL) protocol to ensure millisecond level alignment across data streams and participants. ECG data were streamed in real time from the Callibri sensors using a custom Python script integrated into the LSL environment. To mark experimental events, triggers were sent via the MatNIC toolkit (Neuroelectrics, Barcelona, Spain) to the Neuroelectrics Instrument Controller (NIC) software. Custom-written scripts were used to present the experimental task within the PsychoPy toolbox for Python^42^.

### 2.4 Data preprocessing and metric extraction

#### 2.4.1 Cardiac signal processing and synchronization metrics

Raw ECG signals were first subjected to a narrow bandpass filter (7–13 Hz, 2nd-order Butterworth) specifically designed to isolate the QRS complex while suppressing baseline and high-frequency electromyographic noise. R-peak detection was performed using the algorithm developed by Elgendi and colleagues^43^. The inter-beat interval (IBI) was calculated as the temporal difference between consecutive R-peaks. To address physiological artifacts or signal dropouts, a validation threshold was applied: intervals outside the 0.45-1.2 s range (corresponding to 50-133 beats per minute) were flagged as artifacts. The integrity of each trial was assessed using a 10% artifact-load rule; any trial where more than 10% of the IBIs required correction was excluded from subsequent analysis. Missing or flagged intervals were corrected using linear interpolation and a 5-point moving median window. To facilitate dyadic synchronization analysis, the non-uniformly sampled IBI data were transformed into a continuous timeseries. A cubic spline interpolation was applied to the corrected IBI values, which were then resampled to a uniform frequency of 4 Hz. Finally, to eliminate slow physiological drifts and control for individual differences in baseline heart rate, the resampled signals were detrended to remove linear non-stationarities and *z*-score normalized within each trial.

To quantify physiological synchronization between performers and listeners, we analyzed the processed IBI timeseries. We calculated three metrics of dyadic synchronization using a sliding window approach (window size = 20 s, step size = 1 s):

1. **Sliding window cross-correlation:** Pearson correlations (*r*) were calculated for each window. To account for potential physiological delays between participants, we utilized a cross-correlation method with a maximum lag of ± 5 s, extracting the maximum absolute correlation for each window. Correlation values were Fisher *z*-transformed.
2. **Phase-locking value (PLV):** To assess phase synchrony, we applied a Hilbert transform to the IBI timeseries. The instantaneous phases were extracted, and the PLV was calculated for each window as the magnitude of the mean phase difference vector between performer and listener.
3. **PLV instability (SD PLV):** To evaluate the temporal volatility of cardiac synchronization, we calculated the standard deviation (*SD*) of the PLV across all sliding windows within a given trial. A lower SD PLV indicates a highly stable phase relationship over time, whereas a higher SD PLV reflects an unstable synchronization state.

#### 2.4.2 Continuous and lexical emotion metrics

- **Linear trend in listener’s arousal:** To investigate whether the temporal trajectory of a listener’s arousal is associated with dyadic emotional alignment, we modeled the continuous slider ratings using linear regression over a normalized time axis (0-1). The resulting beta coefficient (slope) quantifies the rate of steady buildup or decay in arousal throughout the duration of the trial.
- **Semantic validation:** Following each trial, participants selected three emotion labels from a predefined list. To validate the label selection task, we extracted affective semantic coordinates (Valence and Arousal) for each selected word triplet using the MEmoLon database^44^, which projects lexical items into a multidimensional affective space. For each trial, a mean semantic score was computed by averaging the coordinates of the three selected words. These scores served as dependent variables in linear mixed models (LMMs) to confirm their alignment with the continuous Likert ratings.
- **GEMS super-factors:** Emotional labels selected by participants were mapped onto the three GEMS super-factors: Sublimity, Vitality, and Unease^33^. For each trial, a prevalence score was calculated for each super-factor by determining the proportion of selected labels belonging to that category relative to the total number of labels chosen (e.g., a triplet containing two Sublimity words and one Vitality word would yield scores of 0.67 and 0.33, respectively).
- **Unique label count:** Operationalized as the total number of unique emotion labels selected by a given participant across the four trials of a specific instruction condition.
- **Word spread:** To measure semantic dispersion, we calculated the arithmetic mean of the Euclidean distances between all possible pairwise combinations of the unique emotion word vectors within the Valence-Arousal-Dominance coordinate space. Higher values indicate the activation of disparate emotional themes (low conceptual coherence), whereas lower values reflect high conceptual coherence.

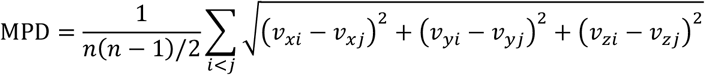

### 2.5 Statistical analyses

All statistical analyses were performed in R (v4.3.0). Figures were generated in R and exported for editing and visual polishing in Adobe Illustrator (Version 27.2; Adobe Inc., San Jose, CA).

#### 2.5.1 Validation of instruction conditions

To evaluate inter-rater reliability among the 7 expert judges, we calculated intraclass correlation (ICC) coefficients. We observed a reasonable agreement across all three evaluation scales: *creativity* (ICC = 0.750, *95% CI* [0.38, 0.93]), *unconventionality* (ICC = 0.633; CI: 0.09-0.90), and *conventionality* (ICC = 0.545, *95% CI* [-0.12, 0.88]). For each rating dimension (conventionality, unconventionality, and creativity), raw scores were z-scored within each rater to account for individual differences in scale usage. These standardized ratings were then averaged across raters. To assess whether the instructions were successfully reflected in the musical output, expert ratings were analyzed using a two-way factorial ANOVA with instruction (Conventional, Creative, Unconventional) and rating metric (Conventionality, Creativity, Unconventionality) as factors. Post-hoc comparisons were performed using Tukey-corrected pairwise tests.

#### 2.5.2 Linear mixed-effects models

We implemented linear mixed-effects models (LMMs) using the lme4 and lmerTest packages. To account for the nested structure of repeated measures within dyads, dyad was specified as a random intercept across all models. *P*-values were adjusted using the False Discovery Rate (FDR) correction to account for multiple comparisons.

Two orthogonal contrasts were applied to the Instruction variable: C_1_ (Creativity): Compares the Creative trials (+2/3) against the mean of Conventional and Unconventional (-1/3 each); and C_2_ (Unconventionality): Evaluates a linear contrast comparing Conventional (-0.5), Creative (0), and Unconventional (+0.5) trials.

Interpersonal familiarity was sum-coded by designating friends as +0.5 and strangers as -0.5.

##### 2.5.2.1 Hypothesis 1: Dyadic emotional alignment and cardiac synchronization

###### Dyadic emotional alignment

We operationalized dyadic emotional alignment as the negative absolute difference between Performer Felt and Listener Felt ratings. This alignment score was calculated separately for the valence and arousal dimensions as follows:

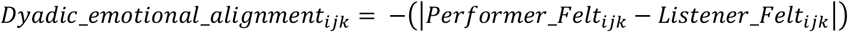

where *i, j*, and *k* represent the specific observation, trial, and dyad, respectively. This transformation ensures that higher values represent greater emotional alignment, spanning a range from -8 to 0, where zero indicates perfect alignment. The following LMM was fitted to evaluate the primary predictors of alignment:

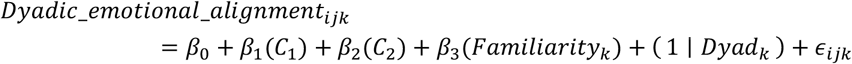

###### Cardiac synchronization

To determine whether emotional alignment facilitates physiological coupling, we examined three cardiac synchronization metrics as dependent variables: phase alignment strength (measured via the phase-locking value, PLV), temporal instability (measured via the standard deviation of PLV, SD PLV), and temporal coupling (measured via the sliding window cross-correlation). The full fixed-effects structure incorporated the condition contrasts (C_1_, C_2_), interpersonal familiarity, and *z*-scored versions of both dyadic valence alignment and dyadic arousal alignment:

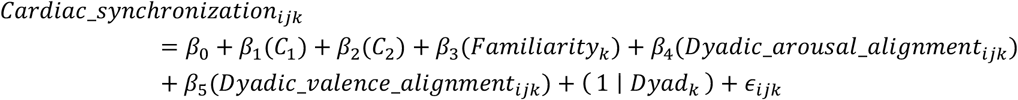

To identify the most parsimonious model architecture, we implemented a stepwise backward elimination procedure (using the lmerTest::step infrastructure). Fixed effects were iteratively removed from the full model one by one based on Satterthwaite approximations for *p-*values, terminating when only statistically significant predictors (*α* = 0.050) remained. The final parameter tables were then subjected to FDR corrections.

###### Intrapersonal emotional alignment

We conducted an exploratory analysis incorporating intrapersonal alignment, defined as the negative absolute difference between an individual’s Expressed and Felt ratings. This was calculated separately for the performer and the listener:

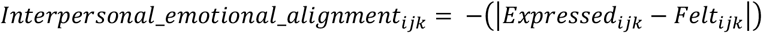

We modeled dyadic emotional alignment and cardiac PLV as dependent variables, with performer and listener intrapersonal alignment included as fixed effects.

###### Linear trend in listener’s arousal

To test if the trajectory of a listener’s arousal shapes alignment, we modeled dyadic emotional alignment separately for arousal and valence. The fixed-effects structure included the instruction contrasts (C_1_, C_2_), Interpersonal Familiarity (Friends vs. Strangers), and the listener’s arousal trajectory (Linear Trend), alongside the interaction terms between the Linear Trend and the instruction contrasts:

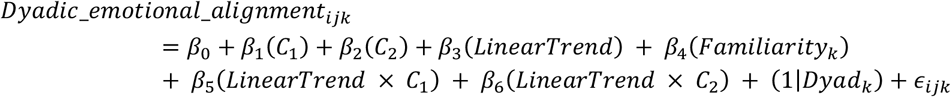

Note that for all equations, *i, j*, and *k* denote the specific observation, trial, and dyad, respectively, and *ε*_*ijk*_ represents the residual error term. The significant predictors of dyadic emotional alignment resulting from the analyses are summarized in a conceptual diagram in Figure 2A.

**Figure 2.**
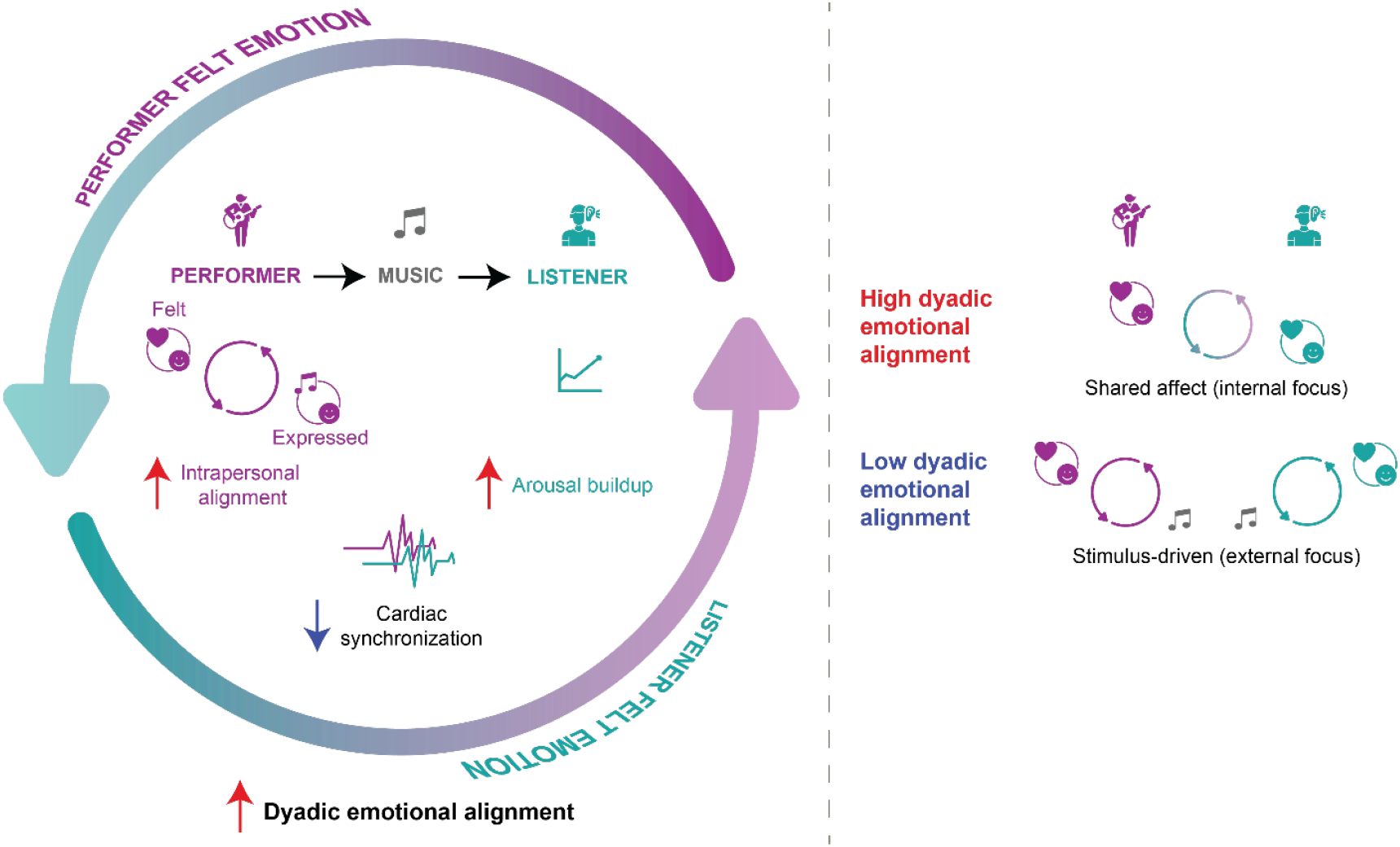
Empirically derived conceptual model of dyadic alignment in live music performance. This schematic summarizes the psychological and physiological mechanisms observed in the current study. **Left:** The circular diagram illustrates the continuous feedback between performer and listener: high intrapersonal alignment in the performer (coherence between felt and expressed emotion) facilitates dyadic emotional alignment, while the listener’s experience is marked by a gradual arousal buildup. Notably, cardiac synchronization is inversely related to dyadic emotional alignment, suggesting a dissociation between subjective and physiological coupling. **Right:** A comparison of alignment states clarifies the hypothesized mechanism. High dyadic emotional alignment is characterized by shared affect between partners (internal focus), whereas low dyadic emotional alignment reflects a state of external anchoring to the musical stimulus. In the latter, both individuals independently track the external musical structure, which accounts for the higher cardiac synchronization observed in the absence of strong shared emotion.

##### 2.5.2.2 Hypothesis 2: Aesthetic experience and semantic metrics

To investigate how instructions shaped aesthetic and lexical experience, we fitted LMMs for each GEMS super-factor (Sublimity, Vitality, Unease) and lexical metric (Unique Label Count, Word Spread). We included Perspective (Felt vs. Expressed) as fixed factor, and Arousal and Valence (z-scored) as continuous covariates. The LMM structure was specified as follows:

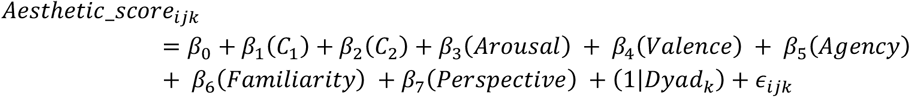

where *i, j*, and *k* represent the specific observation, trial, and dyad, respectively.

## Data and Code Availability

The source data and custom analysis scripts used for the figures generated in this paper have been deposited to the Open Science Framework (OSF) and can be accessed at https://osf.io/fd29p/.

## 3. Results

### 3.1 Validation of instruction conditions

Analysis of the expert ratings confirmed the validity of the experimental design, demonstrating that the three instructions resulted in distinct improvisational profiles (Supplementary Note 3). A two-way factorial ANOVA revealed a significant interaction between instruction and rating (*F*(4,8934) = 680.80, *p* < 0.001). Post-hoc comparisons showed that, for the Conventionality rating, all conditions were significantly differentiated, with the Conventional instruction yielding significantly higher scores than the Creative (*M*_*diff*_ = 0.46, *p* < 0.001) and the Unconventional instruction (*M*_*diff*_ = 1.37, *p* < 0.001). Similarly, for the Unconventionality rating, the Unconventional instruction yielded higher scores than both Creative (*M*_*diff*_ = 0.92, *p* < 0.001) and Conventional (*M*_*diff*_ = 1.40, *p* < 0.001). For the Creativity rating, both the Conventional and Creative instructions elicited significantly higher creativity ratings than the Unconventional instruction (both *p* < 0.001), but did not significantly differ from each other (*M*_*diff*_ = 0.01, *p* = 0.959).

### 3.2 Hypothesis 1: Creativity increases dyadic emotional alignment and cardiac synchronization

#### 3.2.1 Dyadic emotional alignment

To evaluate how different instruction conditions influenced performer-listener emotional alignment, we implemented two orthogonal contrasts in our LMMs. The first contrast (C_1_: Creativity) tested whether instructions to be creative led to a shift in alignment compared to the other two conditions (Creative was coded as +2/3; Conventional and Unconventional were coded as -1/3 each). The second contrast (C_2_: Unconventionality) evaluated the linear effect of increasing unconventionality (Conventional was coded as -0.5, Creative as 0, and Unconventional as +0.5).

Analysis revealed that creative instructions significantly enhanced dyadic emotional alignment. Specifically, the Creativity contrast (C_1_) was significant for both arousal (*b* = 0.385, *95% CI* [0.11, 0.67], *t*(405) = 2.700, *p*_*FDR*_ = 0.022) and valence (*b* = 0.446, *95% CI* [0.14, 0.76], *t*(405) = 2.830, *p*_*FDR*_ = 0.022). The positive coefficients indicate that creative trials fostered higher overall dyadic emotional alignment than the mean of the other two conditions.

In contrast, the Unconventionality contrast (C_2_) revealed a divergent pattern between the two emotion dimensions. While the contrast was significant for valence (*b* = -0.446, *95% CI* [-0.80, - 0.09], *t*(405) = -2.450, *p*_*FDR*_ = 0.029), it was not significant for arousal (*p*_*FDR*_ = 0.440). This indicates that as musical improvisations became increasingly unconventional, valence alignment progressively decreased, whereas alignment along the arousal dimension remained unaffected (Figure 3A). Finally, interpersonal familiarity did not emerge as a significant predictor for either arousal (*p*_*FDR*_ = 0.660) or valence (*p*_*FDR*_ = 0.094).

**Figure 3.**
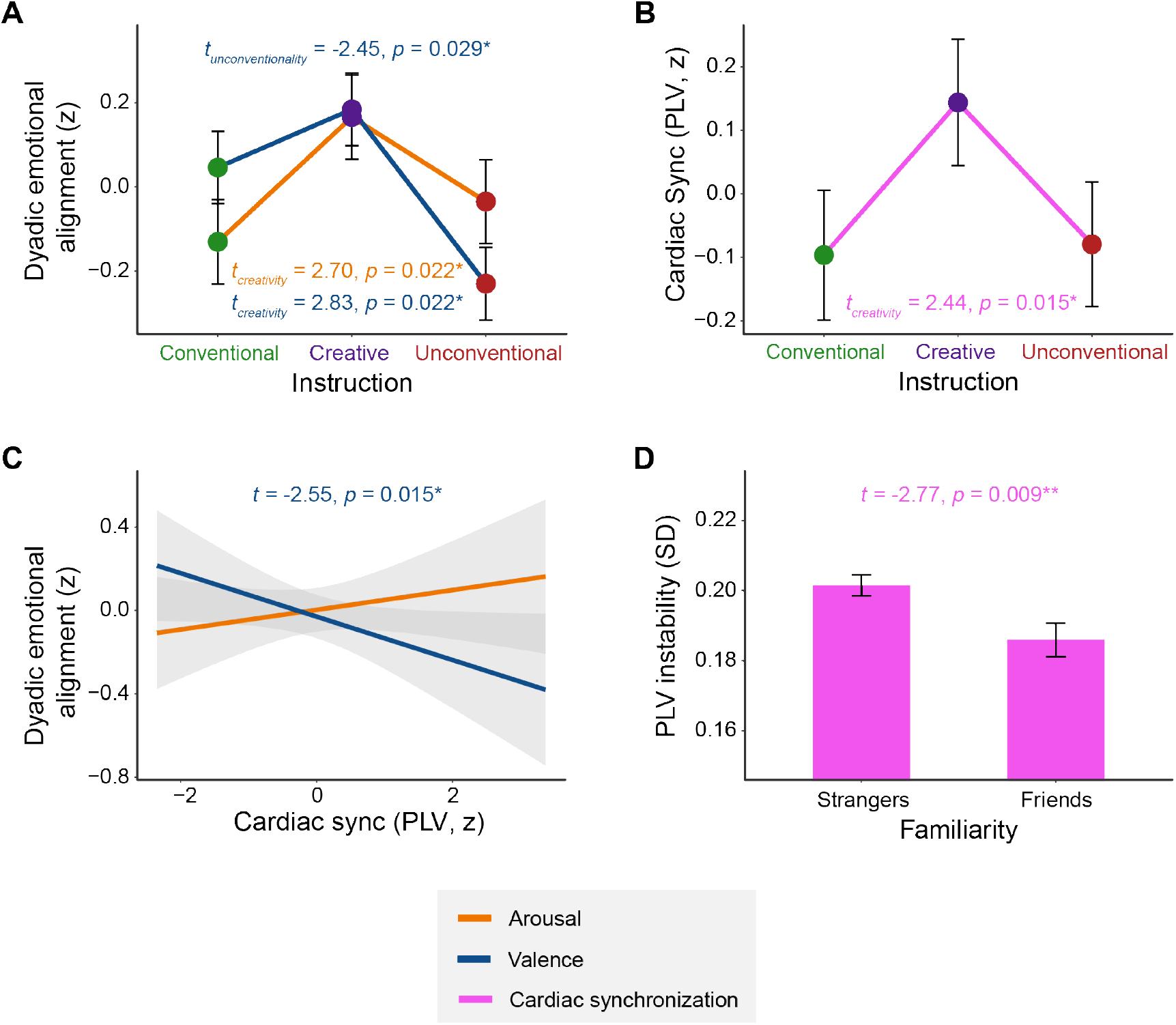
Dyadic emotional alignment and cardiac synchronization. **A** Dyadic emotional alignment scores for the arousal (orange) and valence (blue) dimensions across the three instruction conditions. Statistical values (*t* and *p*) denote the outcomes of the Creativity (C_1_) and Unconventionality (C_2_) contrasts. **B** Cardiac synchronization, measured via the phase-locking value (PLV, pink), across the three instructions. Statistics denote the significant effect of the Creativity contrast. **C** Relationship between PLV and dyadic emotional alignment, separately for arousal and valence. Statistics denote the significant negative slope for dyadic valence alignment (blue). **D** Relationship between the performers’ intrapersonal alignment (the absolute difference between expressed and felt emotions) and PLV. Error bars represent ± 1 standard error of the mean (SEM); shaded areas represent 95% confidence intervals. \**p* < 0.050, \*\**p* < 0.010, \*\*\**p* < 0.001. All *p*-values are FDR corrected.

#### 3.2.2 Cardiac synchronization

To evaluate the relationship between shared affect and physiological coordination, we analyzed metrics of cardiac synchronization as dependent variables. Following a stepwise backward elimination procedure, the Creativity contrast emerged as a significant positive predictor of cardiac phase alignment, measured via the phase-locking value (PLV; *b* = 0.028, *95% CI* [0.00, 0.05], *t*(324) = 2.44, *p*_*FDR*_ = 0.015; Figure 3B). Surprisingly, the model revealed a significant negative effect of dyadic valence alignment on PLV (*b* = -0.014, *95% CI* [-0.03, -0.00], *t*(349) = -2.55, *p*_*FDR*_ = 0.015; Figure 3C), revealing an inverse relationship between valence alignment and PLV. This is explored further in section 3.2.3.

Interpersonal familiarity was a significant predictor of PLV instability (SD PLV) (*b* = -0.016, *95% CI* [-0.03, -0.00], *t*(35.7) = -2.77, *p*_*FDR*_ = 0.009). The negative coefficient demonstrates that friend dyads maintained a more stable cardiac synchronization trajectory over time (i.e., lower volatility) than stranger dyads (Figure 3D). No variables survived model reduction for sliding window cross-correlation (*p* > 0.050).

#### 3.2.3 Exploratory analysis: The role of intrapersonal alignment in dyadic coordination

To further elucidate the previously observed dissociation between dyadic emotional alignment and cardiac synchronization (i.e. the inverse relationship between dyadic valence alignment and PLV), we conducted exploratory analyses to test whether intrapersonal alignment, defined as the convergence between an individual’s felt and expressed emotions, accounted for variability in both metrics.

In the valence domain, a dissociation was observed specifically for the performer: higher intrapersonal alignment significantly predicted increased dyadic alignment (*b* = 0.367, *95% CI* [0.22, 0.51], *t*(431) = 4.890, *p*_*FDR*_ < 0.001; Figure 4A), yet it significantly predicted decreased PLV (*b* = -0.019, *95% CI* [-0.03, -0.01], *t*(349) = -3.350, *p*_*FDR*_ = 0.003; Figure 4B).

**Figure 4.**
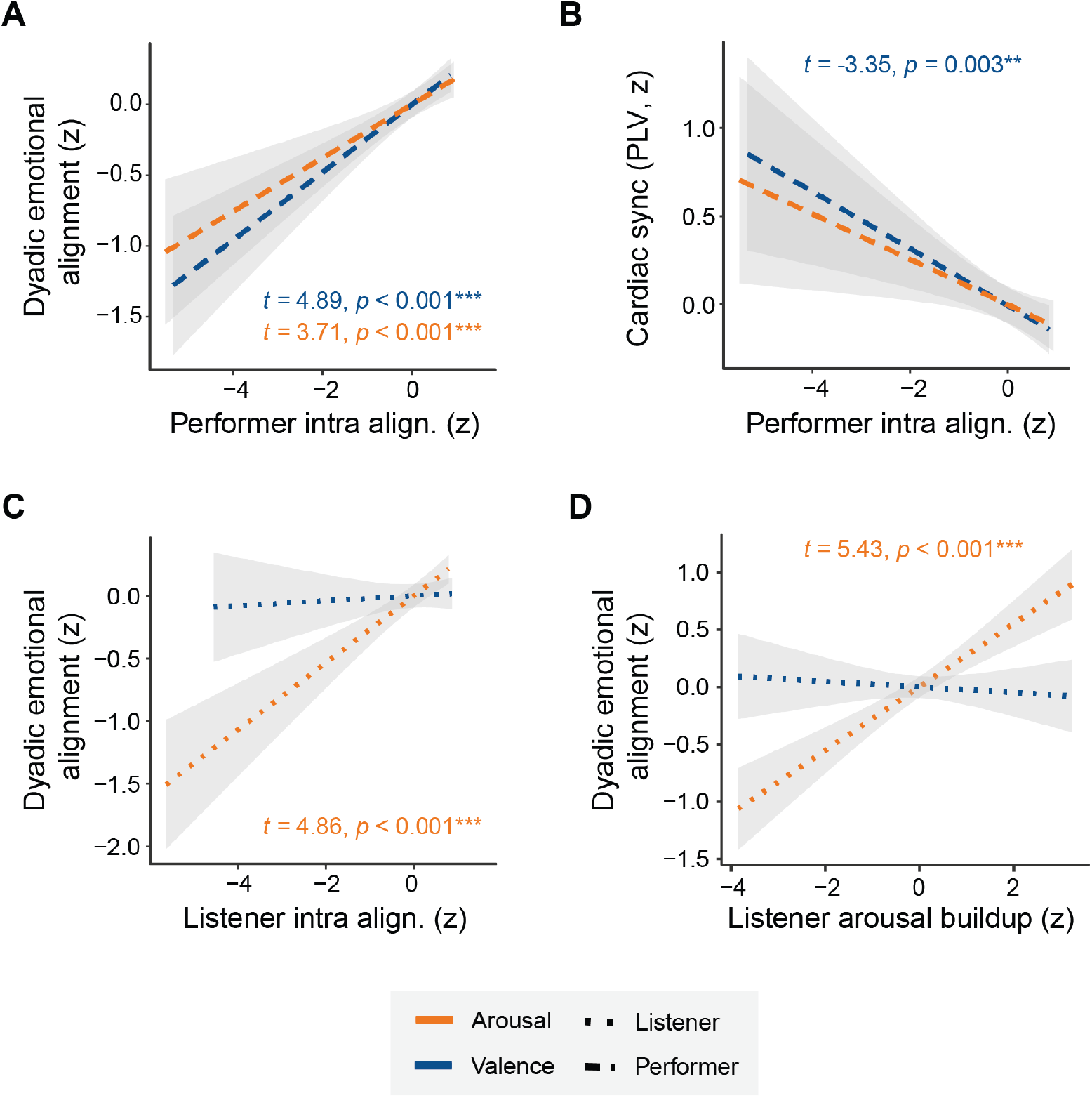
The effect of intrapersonal alignment and listener arousal buildup on dyadic emotional alignment and cardiac synchronization. **A** Relationship between the performer’s intrapersonal alignment (the absolute convergence between felt and expressed emotions) and dyadic emotional alignment, separately for arousal (orange) and valence (blue). **B** Relationship between the performer’s intrapersonal alignment and cardiac synchronization (phase locking value, PLV). **C** Same as **A** but for listeners. **D** Relationship between the listener’s linear arousal buildup and dyadic emotional alignment. Shaded areas represent 95% confidence intervals. \*\**p* < 0.010, \*\*\**p* < 0.001. All *p*-values are FDR corrected.

In the arousal domain, dyadic emotional alignment was positively associated with the intrapersonal alignment of both the performer (*b* = 0.266, *95% CI* [0.13, 0.41], *t*(431) = 3.710, *p*_*FDR*_ < 0.001) and the listener (*b* = 0.348, *95% CI* [0.21, 0.49], *t*(427) = 4.860, *p*_*FDR*_ < 0.001; Figure 4A, C). Crucially, the Creativity contrast remained a significant positive predictor of PLV (*b* = 0.029, *95% CI* [0.01, 0.05], *t*(325) = 2.500, *p*_*FDR*_ = 0.026) even when controlling for intrapersonal alignment. Taken together, this pattern suggests that internal affective consistency within individuals relates differentially, and at times contrastingly, to emotional versus physiological coordination across dyads.

#### 3.2.4 Linear trend of listener arousal

Analysis of listeners’ continuous arousal ratings revealed that the linear trajectory of listener arousal was a significant predictor of dyadic arousal alignment (*b* = 0.380, *95% CI* [0.24, 0.52], *t*(435) = 5.430, *p*_*FDR*_ < 0.001). This indicates that as listeners’ arousal progressively increased over the course of an improvisation, the emotional alignment between the performer and listener strengthened. In contrast, dyadic valence alignment was not significantly predicted by this linear increase in listener arousal (*p*_*FDR*_ = 0.740; Figure 4D).

To determine if instruction modulated this relationship, we evaluated the interaction terms between the linear arousal trajectory and the instruction contrasts (C_1_, C_2_). For both arousal and valence alignment models, no significant interactions emerged (all *p*_*FDR*_ > 0.060). This suggests that the positive relationship between a listener’s arousal trajectory and dyadic emotional alignment remained relatively invariant across the different instruction conditions.

### 3.3 Hypothesis 2: Aesthetic quality and semantic richness

#### Qualitative thematic profiles

To characterize the qualitative themes of emotional states underlying dyadic interaction, we examined the distribution of discrete emotion labels across the different instruction conditions (Table 1). Conventional improvisations were dominated by low-arousal positive labels (e.g., Relaxation, Pleasure). Creative improvisations shifted the emotional landscape toward states associated with openness and aesthetic elevation (e.g., Freedom for performers, Tenderness and Dreaminess for listeners). Unconventional improvisations were characterized by high-arousal, negatively valenced labels (e.g., Tension, Aggression, Rebelliousness). See Supplementary Note 4 for word cloud visualizations.

**Table 1.** Top 3 emotion labels separated by instruction and agency. Values represent percentage of trials in which a label was selected within participants’ top-3 lists, aggregated across felt and expressed perspectives.

|  | Rank | Conventional | Creative | Unconventional |
| --- | --- | --- | --- | --- |
| Performer | 1 | Relaxation (29.7%) | Freedom (21.6%) | Energy (36.5%) |
|  | 2 | Pleasure (23.3%) | Tension (18.2%) | Tension (35.1%) |
|  | 3 | Amusement (21.6%) | Relaxation (17.9%) | Aggression (23.0%) |
| Listener | 1 | Relaxation (26.0%) | Relaxation (20.9%) | Tension (34.1%) |
|  | 2 | Pleasure (18.9%) | Tenderness (16.6%) | Energy (22.3%) |
|  | 3 | Happiness (17.6%) | Dreaminess (15.9%) | Aggression (18.2%) |

#### Validation of the label selection task

To ensure that the discrete emotion labels provided a valid index of the underlying emotions, we validated the semantic coordinates of the selected word triplets against the Likert ratings. LMMs confirmed a significant relationship between Likert ratings and the chosen semantic vectors for both Arousal (*b* = 0.07, *95% CI* [0.06, 0.09], *t*(1739.0) = 12.76, *p*_*FDR*_ < 0.001) and Valence (*b* = 0.48, *95% CI* [0.44, 0.52], *t*(1739.0) = 24.64, *p*_*FDR*_ < 0.001). This confirms that self-reported Likert ratings consistently mapped onto the semantic properties of the selected words, establishing a reliable link between the quantitative scales and qualitative descriptors.

#### Aesthetic quality

We fitted LMMs predicting each GEMS super-factors with instruction (creativity and unconventionality contrasts), agency (performer vs. listener), perspective (felt vs. expressed), and interpersonal familiarity (friends vs. strangers) as fixed effects and valence and arousal as covariates. Dyad was specified as a random intercept. The Creativity contrast was modeled as a planned contrast testing the deviation against the mean of the other two conditions (Creative coded as +2/3 and Conventional/Unconventional as -1/3).

Creative improvisations were associated with higher Sublimity (*b* = 0.275, *95% CI* [0.19, 0.36], *t*(1739) = 6.020, *p*_*FDR*_ < 0.001), and reduced Vitality (*b* = -0.158, *95% CI* [-0.23, -0.09], *t*(1739) = - 4.45, *p*_*FDR*_ < 0.001) and Unease (*b* = -0.130, *95% CI* [-0.21, -0.05], *t*(1739) = -3.270, *p*_*FDR*_ = 0.002; Figure 5A).

**Figure 5.**
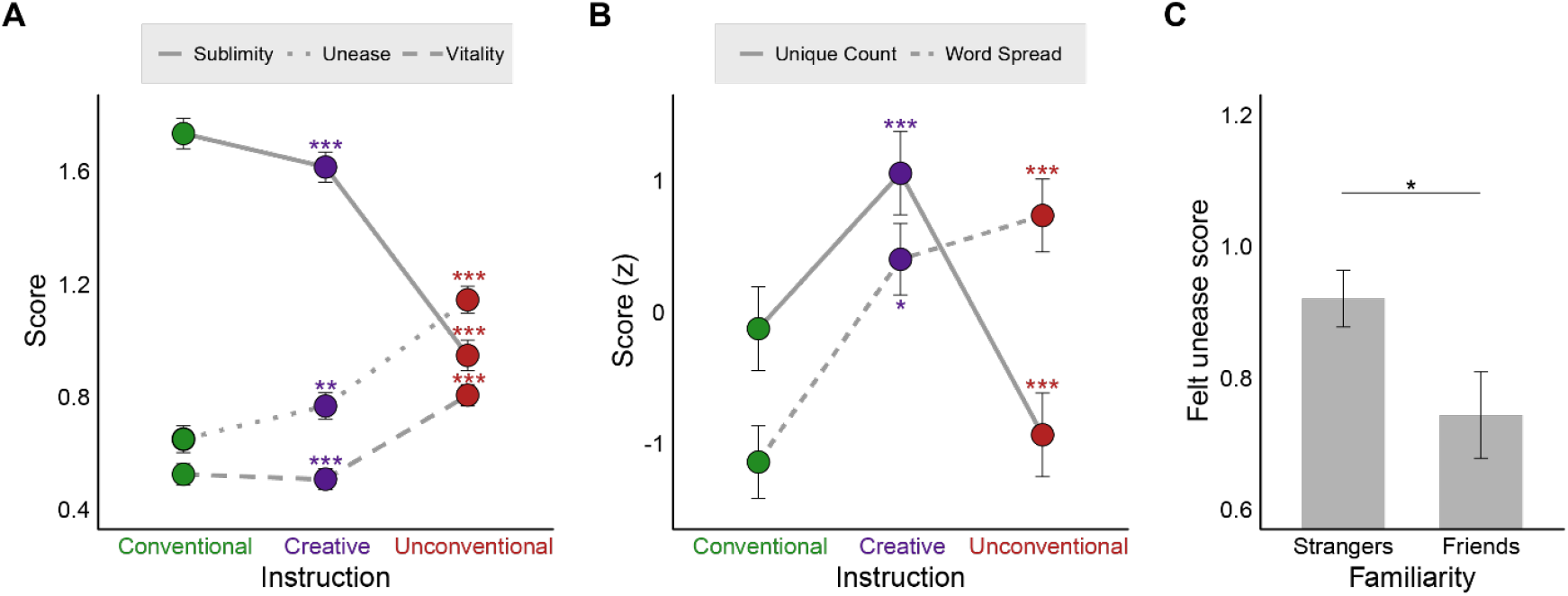
Aesthetic qualities, semantic richness, and interpersonal familiarity. **A** Estimated marginal means (EMMs) for the GEMS dimensions (Sublimity, Vitality, Unease) across the three instruction conditions. **B** EMMs of the standardized semantic richness metrics (Unique emotion label count and Semantic word spread) across instructions. **C** Main effect of interpersonal familiarity on felt Unease. Error bars represent ±1 SE. Statistical annotations denote the FDR-corrected *p*-values and *t*-statistics for the significant contrasts.

The Unconventionality contrast was modeled as a linear contrast reflecting a progressive shift from conventional through creative to unconventional improvisation (Conventional = -0.5, Creative = 0, Unconventional = +0.5). The Unconventionality contrast showed the opposite pattern, with decreased Sublimity (*b* = -0.787, *95% CI* [-0.90, -0.68], t(1739) = -14.100, *p*_*FDR*_ < 0.001), and increased Unease (*b* = 0.495, *95% CI* [0.40, 0.59], *t*(1739) = 10.200, *p*_*FDR*_ < 0.001) and Vitality (*b* = 0.281, *95% CI* [0.20, 0.37], *t*(1739) = 6.480, *p*_*FDR*_ < 0.001; Figure 5A). Additionally, friends reported lower felt Unease than strangers (*b* = -0.177, *95% CI* [-0.33, -0.02], *t*(37) = -2.330, *p*_*FDR*_ = 0.046; Figure 5C). Neither agency nor perspective yielded significant effects (all *p*_*FDR*_ > 0.200).

#### Semantic richness and spread

Beyond categorical affect, we examined how instructions shaped the complexity of the shared emotional representation space, indexed by unique label diversity and semantic dispersion. Creative trials increased both the number of unique emotion labels selected (*b* = 0.621, *95% CI* [0.49, 0.76], *t*(1739) = 9.120, *p*_*FDR*_ < 0.001) and their semantic dispersion (*b* = 0.043, *95% CI* [0.01, 0.07], *t*(1739) = 2.790, *p*_*FDR*_ = 0.011), indicating a broader and less constrained emotional space.

Conversely, increasing Unconventionality significantly reduced unique label diversity (*b* = -0.316, *95% CI* [-0.48, -0.15], *t*(1739) = -3.800, *p*_*FDR*_ < 0.001), but expanded semantic spread (*b* = 0.133, *95% CI* [0.10, 0.17], *t*(1739) = 7.070, *p*_*FDR*_ < 0.001). Finally, performers exhibited greater unique label diversity than listeners (*b* = 0.432, *95% CI* [0.31, 0.56], *t*(1739) = 6.750, *p*_*FDR*_ < 0.001; Figure 5B).

## 4. Discussion

In this study, we investigated whether creativity shapes the alignment of emotional and physiological states between performers and listeners during live music improvisation. We observed that (1) Creative improvisation heightened both dyadic emotional alignment and cardiac phase synchronization between partners; (2) Dyadic emotional alignment was dependent on both partners, with higher performer intrapersonal alignment and a progressive linear increase in listener arousal both predicting greater dyadic emotional alignment; and (3) Creative improvisations elicited a distinct aesthetic profile characterized by elevated sublimity, low unease, and reduced vitality, and were associated with greater semantic richness and dispersion in the selected emotional descriptors.

### Creativity enhances performer-listener emotional alignment and cardiac synchronization

Creative instructions significantly enhanced both dyadic emotional alignment and cardiac phase synchronization. This effect was grounded in a genuine manipulation of performance style: blind expert ratings confirmed that the three different creative instruction conditions produced highly recognizable, differentiated improvisational profiles. Importantly, the observed dyadic emotional alignment reflects convergence specifically in felt rather than merely expressed emotion, a critical empirical distinction that moves the field beyond prior work focused primarily on emotion recognition.

At the physiological level, creative improvisations were associated with increased cardiac phase synchronization (PLV), indicating a robust temporal coupling of autonomic activity. These results are in line with concert work demonstrating that audience members synchronize their cardiac and autonomic channels during shared musical exposure, and that heightened attentional engagement during listening predicts stronger physiological synchrony^45^. Furthermore, emerging evidence from live performance contexts indicates that autonomic synchrony is modulated by the musical piece itself^7^, consistent with the idea that specific temporal-dynamic properties of an auditory stimulus entrain physiological timing^10,45^. In our study, the instruction to “be creative” may have driven performers to generate improvisations that amplify such structural, attention-gaining features, thereby concurrently elevating subjective emotional alignment and cardiac coupling.

Performer-listener coupling during live improvisation likely emerges from multiple sources simultaneously: entrainment to shared musical timing, shared attention, and covert respiration changes driven by shared arousal dynamics. Indeed, in interactive dyads vocalizing together, heart rate variability coupling increases during sustained, synchronous vocalizations; an effect largely mediated by respiratory sinus arrhythmia driven by task-constrained breathing patterns^46^. Similarly, in Javanese gamelan performance, physiological synchronization is positively associated with self-reported shared flow states during improvised playing, yet negatively associated during traditional playing, suggesting that the act of improvisation strengthens the link between physiological coupling and shared experiential states^47^. Autonomic synchrony emerges within shared affective and interactive contexts^7,10^. It is therefore highly plausible that creative intent enhances the improvisational features of a performance, intensifying the performer-listener interaction and driving dyadic alignment. Yet, the relationship between emotional alignment and physiological synchronization proved more complex than anticipated, as not all results fit a simple account.

Contrary to expectations, higher dyadic alignment along the valence dimension was associated with lower cardiac phase synchronization, revealing a striking dissociation between subjective emotional convergence and autonomic coupling. This dissociation suggests that subjective and physiological coupling are supported by partially distinct modes of engagement, in contrast to prior evidence suggesting that when individuals evaluate a narrative performance similarly, their autonomic dynamics become more temporally aligned^17^. As a post-doc explanatory hypothesis consistent with our data, we propose that high dyadic emotional alignment reflects a state of “internal focus”. Under this account, partners may become highly absorbed in their internal affective state space and less driven to track external, moment-to-moment physical anchors, thereby reducing stimulus-locked physiological synchrony. Indeed, evidence from narrative paradigms demonstrates that when a participant’s attention is diverted inward (e.g., toward an internally demanding or distracting cognitive task), inter-subject physiological synchrony is attenuated^48^.

Conversely, when dyadic emotional alignment is low, dyads may rely more on tracking shared, external sensorimotor anchors (such as tempo, dynamics, or visual performer movement cues), yielding stronger cardiac phase synchronization despite their divergent emotional interpretations. Indeed, tracking of external physical cues can robustly drive cardiac synchronization in the absence of shared emotion^20,48^. For instance, during movie viewing, higher heart rate synchronization correlates with superior memory for content, and this synchronization strength changes with attentional states (active attention vs. distraction)^20^. Similarly, physiological synchrony is higher when individuals actively attend to narrative stimuli than when they are distracted or explicitly instructed to focus attention inward on a distracting task^48^.

In our study, the cardiac synchronization observed in the absence of strong emotional alignment may therefore reflect a state of shared entrainment to common stimulus dynamics and audiovisual performance cues, representing a state of co-attunement to external features rather than a true convergence of inner emotional experience. Crucially, this dissociation does not contradict the finding that creative instructions simultaneously enhance both emotional alignment and cardiac synchronization. Rather, creativity appears to operate as a global driver that raises the baseline of both metrics concurrently. In contrast, the inverse relationship between emotional alignment and cardiac synchronization reflects a finer-grained attentional dynamic operating within that context: as the shared emotional experience deepens, attention shifts inward, partially decoupling physiological entrainment from subjective convergence. While this offers a compelling interpretation of the observed PLV-emotional alignment dissociation, it remains a hypothesis that warrants testing in future neuroimaging work. Taken together, these findings demonstrate that while creativity boosts emotional alignment and physiological coupling, these effects are dissociable across subjective and autonomic levels. Future research should investigate how specific acoustic features (e.g., temporal structure, expressive variability, spectral dynamics) interact with visual performance gestures and social dynamics of the live environment to shape this interpersonal coupling.

### Dyadic emotional alignment emerges from performer intrapersonal alignment and listener arousal buildup

Performer-listener emotional alignment was shaped by parallel processes operating within each partner: the closer a performer’s felt and expressed emotions converged, the greater the dyadic alignment with the listener. This effect was robust for performers across both valence and arousal, whereas for listeners it was restricted to arousal, suggesting role-specific contributions to the shared emotional experience.

A performer’s intrapersonal alignment is externalized through their musical output, which in turn shapes listener responses^6,49^. We propose that when performers’ felt and expressed emotions align, the resulting musical signal becomes less ambiguous. This enhanced communicative clarity facilitates more accurate emotion inference on the part of the listener, ultimately reducing the psychological distance between performer and listener. While felt-expressed congruence remains underexplored, multiple lines of work support this framework. For instance, this view is consistent with accounts of musical communication asserting that interpersonal emotional convergence arises from the decoding of expressive cues rather than direct access to a performer’s internal states^50^.

In live performance settings, listeners explicitly link perceived emotions to performers’ expressive actions; dynamic increases in brightness, tempo, and acoustic volume are consistently characterized as driving heightened arousal and positive imagery, whereas hesitancy or timing violations evoke psychological tension and anxiety^51^. These observations support the interpretation that when a performer’s internal affective state is congruent with the expressive decisions they enact, listeners are more likely to emotionally converge with that trajectory. Furthermore, live concert research indicates that shared aesthetic experiences and physiological synchronization scale with the audience’s immersion and state of being “moved”^45^. Consequently, if felt-expressed coherence increases the expressive clarity of a performance, it serves as a primary catalyst for dyadic emotional alignment. Crucially, this heightened absorption in the shared experience may prioritize internal dyadic resonance over the monitoring of external musical cues.

This provides a plausible mechanistic explanation for the underlying dynamics: a performer’s internal emotional authenticity serves as the foundation for successful emotion transmission. When felt and expressed emotions align, the resulting expressive clarity minimizes ambiguity in the acoustic signal, facilitating seamless emotional inference by the listener, ultimately leading to a high degree of dyadic emotional alignment.

Paradoxically, while this felt-expressed alignment increases dyadic emotional alignment, it appears to actively decouple the dyadic cardiac synchronization. This intrapersonal finding mirrors the interpersonal valence dissociation described earlier, providing corroborating evidence for our dual-process framework. This supports our hypothesis that as a performer achieves internal emotional alignment, their cognitive resources are reallocated away from tracking external musical cues shifting their attention inward. Because cardiac synchronization is largely tied to the tracking of external musical structures^8,52^, this internal focus occurs at the expense of external physiological entrainment. Subjective emotional experience and physical entrainment thus represent distinct, potentially competing pathways of dyadic coupling.

Stronger emotional performer-listener alignment was also associated with a gradual linear increase in listener arousal over time. This linear trajectory of arousal can be interpreted as a behavioural signature of sustained attention, which subsequently enhances dyadic alignment. Previous research has shown that physiological synchronization peaks at salient musical events and accumulates across time^8,53^. This finding could also be related to predictive processing accounts, which posit that individuals continuously generate and update expectations during music listening^54–56^.

Within this predictive framework, music listening consists of cycles of curiosity, attention, and positive affect that emerge when prediction errors are resolved faster than anticipated^57^. Furthermore, musical pleasure peaks at intermediate levels of predictability and uncertainty, supporting the idea that engagement is sustained when listeners can successfully track and update an evolving acoustic structure^58–61^. Accordingly, the observed gradual increase in arousal may reflect a constant engagement with the evolving musical structure and the ongoing updating of internal predictions, marking a state of heightened attention during a continuous learning process focused on the reduction of structural uncertainty.

Under this predictive account, an arousal buildup serves as a proxy of the listener’s inference process becoming coupled to the unfolding musical structure. This interpretation is consistent with evidence showing that, compared to standard repertoire performance, musical improvisation elicits greater cognitive engagement in listeners alongside greater relaxation in performers^62^.

Ultimately, these findings support the interpretation that an increasing arousal state enhances the probability that listeners’ responses become temporally aligned with the unfolding musical structure, which in turn promotes dyadic emotional alignment. Taken together, the current literature and our findings suggest that dyadic emotional alignment in live music peaks when two conditions co-occur: (i) the performer provides an unambiguous expressive signal backed by felt-expressed congruence, and (ii) the listener closely tracks the evolving musical structure, culminating in a sustained, progressive linear increase in arousal. An illustration of this proposed mechanism is presented in the conceptual model of Figure 2, offering a foundation for future experimental validation.

### Creative improvisations elicit a distinct aesthetic-emotion profile and increase semantic richness

Creative instructions elicited a distinct aesthetic profile characterized by elevated sublimity and reduced unease and vitality. Within the GEMS framework, sublimity operates as a higher-order factor comprising wonder, transcendence, tenderness, nostalgia, and peacefulness^33^. Prior research emphasizes that aesthetic emotions, such as awe, fascination, and the psychological state of being “moved”, are triggered by the aesthetic appeal of a stimulus^63^. Furthermore, aesthetic judgement is considered a higher-level process that operates over and above low-level perceptual features, relying on criteria that encompass perceived beauty, novelty, expressive skill, and stylistic mastery^64^.

Additionally, musical awe includes a positive form linked to pleasure, joy, surprise, wonder, and tenderness^64^. For instance, in a live opera setting, Balteş and Miu^64^ reported high sublimity, low vitality, and moderate unease. In this view, creative instruction may bias appraisals toward positive emotions (low unease) and high transcendence (high sublimity), rather than toward energetic activation (low vitality). This pattern aligns with previous research on aesthetic emotions in music, which suggests that complex, novel musical structures tend to evoke transcendence and beauty rather than high-arousal activation^34,35^.

In addition, creative instructions produced a broader and more diverse set of emotional descriptors, reflected by an increase in both unique label diversity and semantic spread. This indicates that creativity is associated with a wider exploration of the affective space. Indeed, creative production is known to engage more widely distributed semantic activation, consistent with associative accounts of creativity proposing that creative thought relies on the recruitment of more remote associations within semantic memory^36,39^.

Semantic network approaches have similarly shown that higher creative capacity is associated with more flexible, less clustered semantic structures, enabling cognitive search processes to access more distant conceptual neighbours^38^. Our finding that performers generated a greater number of unique descriptors than listeners is consistent with this interpretation; the generative act of improvisation demands a much broader, active exploration of semantic space than the more constrained task of passive music listening. Ultimately, the finding that creative improvisations increased semantic dispersion demonstrates that creativity evokes highly divergent aesthetic states, rather than a single prototypical emotional category.

## Conclusion

Our findings demonstrate that creative improvisation enhances both dyadic emotional alignment and cardiac phase synchronization within performer-listener dyads. Furthermore, creativity also elicits a distinct aesthetic emotional profile, characterized by elevated sublimity alongside reduced unease and vitality, as well as an increased semantic richness and unique label diversity in the chosen emotional descriptions. We elucidate that this dyadic alignment emerges from complementary mechanisms, specifically involving the performer’s intrapersonal consistency and the listener’s engagement over time. These findings support a dual-process account of interpersonal coupling, linking creative cognition to coordinated subjective emotional and physiological states during live social interaction.

## Acknowledgements

We would like to thank Spyridon Dimitrakakis and Odysseas Chalkias for their contributions to data collection and technical support. We are also grateful to all the performers and listeners who voluntarily participated in this study.

## Author Contributions

I.Z. conceived the study, formulated the research questions, designed the experimental paradigm, programmed the data collection, collected and analyzed the data, and wrote the original draft of the paper. C.G. contributed to the formulation of the research questions and to data analysis both conceptually and by producing analysis scripts, and reviewed and edited the paper. C.D.B.L. contributed to the conceptualization of the study, the development of the experimental paradigm and the formulation of the analytical ideas, and reviewed and edited the paper. C.P. and C.A. contributed to the methodology, contributed to discussions of the findings, provided resources and supervision, and reviewed and edited the paper. All authors approved the final version of the paper.

