## Supplementary Material for "Creativity drives performer–listener emotional and physiological alignment in live music improvisation"

**This file includes:**

Note 1: Questionnaires

Note 2: Emotional assessment – list selection

Note 3: Validation of instruction conditions

Note 4: Word clouds of discrete emotion labels

***Supplementary Note 1: Questionnaires***

The questionnaires administered were the following:

1. *Goldsmiths Musical Sophistication Index (Gold-MSI)*^1^*.* The factor used from this questionnaire is Musical Training (dimension 3), which includes 7 self-report statements regarding typical musical education experience and musical skills (e.g., “I would not consider myself a musician”).
2. *Improvisation Experience Question.* A question was administered to assess the duration of participants’ involvement in musical improvisation. Participants were asked to indicate their experience in years by selecting one of the following categories: 0, 0.5, 1, 2, 3, 4-6, 7-10, or 11 or more years. For statistical analysis, these categorical responses were converted into numeric values using the midpoint of each range (e.g., 5 years for the "4-6" category and 8.5 years for the "7-10" category), while "11 or more" was coded as 12 years.

The translation from English to Greek for questionnaire 1 was performed based on the protocol proposed by Guillemin and colleagues^2^. Specifically, an independent psychologist with a native Greek and a high level of English translated the questions into Greek (forward translation). Subsequently, a native English speaker with a high level of Greek and who had not seen the original version of the questions, retranslated the Greek version into English (back translation) to ensure semantic equivalence and cultural appropriateness.

***Supplementary Note 2: Emotional assessment – list selection***

To investigate qualitative descriptions of the emotional experience beyond arousal and valence, participants selected three labels from a list of 31 music-specific emotion terms. We adopted the Geneva Emotional Music Scale (GEMS)^3^, a theoretically grounded framework designed to capture music-induced emotions. While the core structure followed the original GEMS taxonomy, twelve additional labels were included to better capture interpersonal dynamics (e.g., empathy), aesthetic qualities (e.g., beauty), and agency-related experiences (e.g., freedom). For analysis, selected labels were aggregated into the second-order GEMS factors, Sublimity, Vitality, and Unease, following established conventions. The full mapping between labels and factors is provided in Supplementary Table S1 below.

**Supplementary Table S1. Classification of emotion labels according to GEMS factors.**

| GEMS factor | GEMS dimensions |
| --- | --- |
| Sublimity | Wonder (*Happiness, Beauty, Being moved, Enthusiasm, Sublimity*);  Transcendence (Spirituality, *Freedom*);  Tenderness (Tenderness, Love, Eroticism, *Empathy*);  Nostalgia (Nostalgia, *Dreaminess*);  Peacefulness (Relaxation, Pleasure) |
| Vitality | Power (Strength, Energy, *Triumph*);  Joy (*Amusement*, *Curiosity*) |
| Unease | Tension (Tension, Aggression, Fear, *Rebelliousness, Anger*);  Sadness (Sadness, *Loneliness, Pain, Vulnerability*) |

*Note:* Categories are based on the Geneva Emotional Music Scale^3^. Standard GEMS categories are listed first, followed by the specific labels used in the present study. Labels added for this study are indicated in *italics*.

***Supplementary Note 3: Validation of instruction conditions***

To verify that performers adhered to the experimental instructions, an independent pool of seven expert musicians evaluated all improvisations on three dimensions: conventionality, creativity, and unconventionality. Ratings were z-scored within rater and dimension prior to aggregation. Results are visualized in Supplementary Figure S1 below.

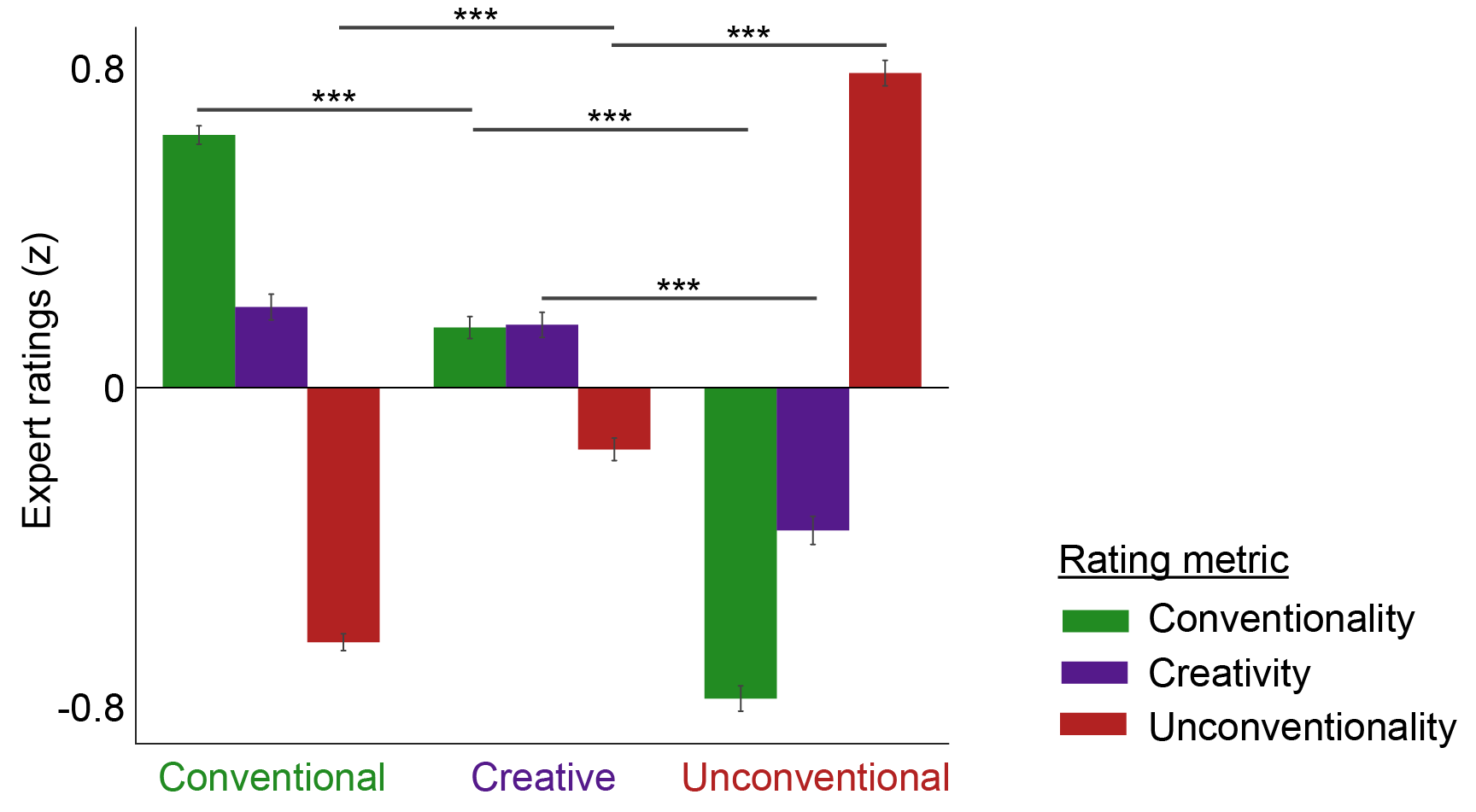

**Supplementary Figure S1. Expert validation of experimental instructions.** Bar chart showing standardized expert ratings (z-scores) across the three instructions (Conventional, Creative, Unconventional) for each rating dimension (Conventionality, Creativity, Unconventionality). Error bars represent ±1 standard error of the mean (SEM). *** *p* < 0.001.

***Supplementary Note 4: Word clouds of discrete emotion labels***

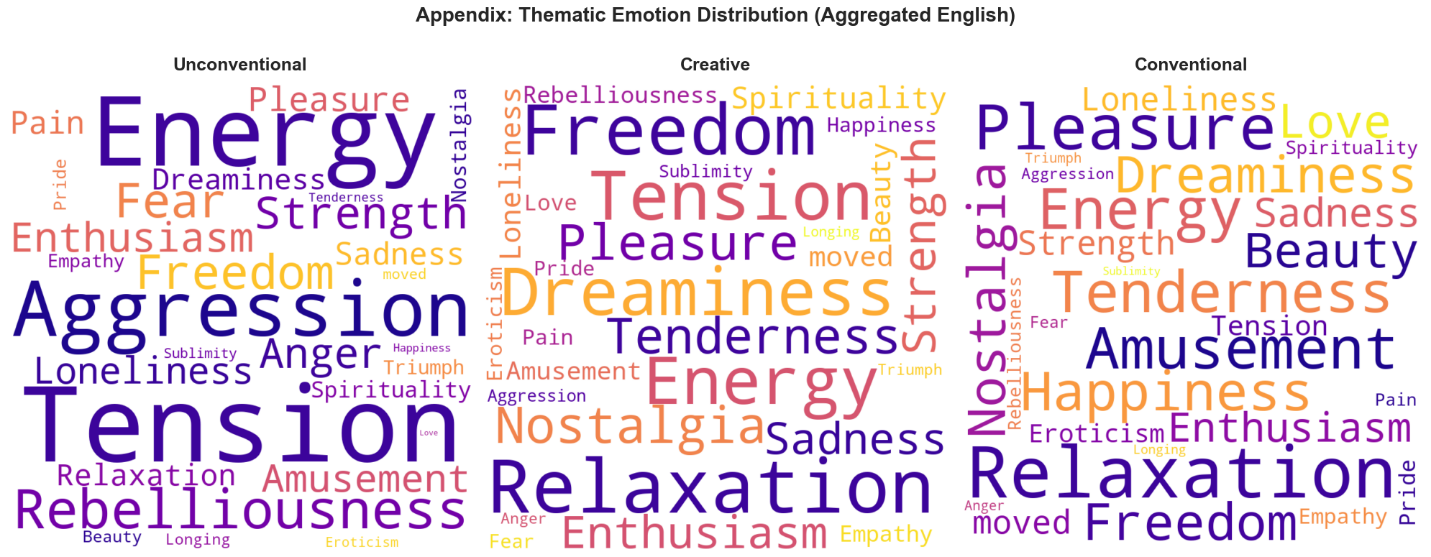

**Supplementary Figure S3. Thematic distribution of emotion labels across musical instructions.** Word clouds represent the relative frequency of discrete emotion labels selected by participants (top-3 selections), aggregated across roles (performer, listener) and perspectives (felt, expressed). Word size reflects selection frequency. All labels were translated into English for consistency. A fixed layout (random_state = 42) was used to facilitate visual comparison across conditions.
